# Interactive dynamics of matrix adhesion and reaction-diffusion predict diverse multiscale strategies of cancer cell invasion

**DOI:** 10.1101/2020.04.14.041632

**Authors:** Durjay Pramanik, Mohit Kumar Jolly, Ramray Bhat

## Abstract

The metastasis of malignant epithelial tumors begins with the egress of transformed cells from the confines of their basement membrane to their surrounding collagenous stroma. Invasion can be morphologically diverse, ranging from dispersed mesenchymal cells to multicellular collectives. When breast cancer cells are cultured within basement membrane-like matrix (BM), or Type 1 collagen, or a combination of both, they exhibit collective-, dispersed mesenchymal-, and hybrid collective-dispersed (multiscale) invasion, respectively. In this paper, we asked how distinct these invasive modes are with respect to the cellular and microenvironmental cues that drive them. A rigorous computational exploration of invasion was performed within an experimentally motivated Cellular Potts-based modeling environment. The model comprises of adhesive interactions between cancer cells, BM- and collagen-like extracellular matrix (ECM), and reaction-diffusion-based remodeling of ECM. The model outputs were parameters cognate to dispersed- and collective- invasion. Input sweeps gave rise to a spatial output distribution that consisted of dispersed-, collective- and multiscale- invasion. K-means clustering of the output distribution followed by silhouette analysis revealed three optimal clusters: one signifying indolent invasion and two representing multiscale invasions, which we call collective-multiscale (CMI), and dispersed multiscale invasion (DMI), respectively. Constructing input-output mapped phenotypic spaces suggested that adhesion to BM- and collagen- matrix specify CMI and DMI respectively. Parameter perturbations confirmed these associations and revealed how the cellular phenotype may transition between the three states. Our systems-level analysis provides quantitative insights into how the diversity in matrix microenvironments may steer invasion into distinct phenotypic modes during metastasis.

## Introduction

The details of the beginnings of cancer progression determine not just the kinetics of its metastasis but also its response to therapeutic efforts (Haeger et al., 2020; Nieto et al., 2016). Epithelia from malignant tumors proliferate and breach their covering laminin-rich basement membrane (BM) barriers to migrate to the surrounding connective tissue consisting of fibroblasts, other resident cells, and extracellular matrix (Nelson and Bissell, 2005; Pally et al., 2019; Pickup et al., 2014). The latter is rich in Type 1 collagen and other fibrillar proteins as well as proteoglycans (Di Lullo et al., 2002; Hynes, 2009).

Bidirectional interactions including adhesion and degradation between cancer cells and their surrounding tissue microenvironment shape the nature of their migration (Anderson, 2005; Nissen et al., 2019) Cancer cell invasion is broadly classified into unicellular or multicellular categories (Roussos et al., 2011). Solitary cancer epithelia disperse and migrate through ECM in a fibroblast-like manner (Madsen and Sahai, 2010). Such a dispersed migration pattern often concurs with a series of changes in gene expression, intercellular adhesion and cell shape, known as the epithelial to mesenchymal transition or EMT (van Zijl et al., 2011). The spindle-shaped ‘mesenchymal’ cells interact extensively with ECM in their process of migration (unlike the other unicellular migratory mode: amoeboid, where the adhesion with matrix is minimal) (Friedl et al., 2004; Gadea et al., 2008; Huang et al., 2014; Pankova et al., 2010; Sanz-Moreno et al., 2008). Collective cell invasion involves migration of cellular ensembles that retain adhesive and communicative contacts with each other through their movement (Cheung et al., 2013; Friedl et al., 2012). These diverse phenotypes are also reflected in blood-borne dissemination of circulating tumor cells as individual cells or multicellular clusters (Aceto et al., 2014; Bocci et al., 2019).

The migratory dynamics of cancer epithelia can also transition among the abovementioned types: dispersed (mesenchymal or amoeboid), and collective; such plasticity may further drive phenotypic heterogeneity and symbiotic behavior in cancer invasion patterns (Hecht et al., 2015; Huang et al., 2015; Lintz et al., 2017). As cancer cells migrate to newer microenvironments, transitions from mesenchymal to epithelial morphologies could appropriately render invasive single cells more adhesive to each other (Krakhmal et al., 2015). Collectively migrating strands of cells are typically thought of as hybrid epithelial/mesenchymal (E/M) phenotype (Nagai et al., 2020); *in vitro, in vivo* and *in silico* evidence for existence, plasticity and aggressiveness of such hybrid E/M cells has been mounting (Jolly et al., 2019). Thus, it is not surprising to identify unique migratory behaviors of neoplastic cells that are phenotypically intermediate between their dispersed and collective counterparts; for instance, multicellular streaming of an amoeboid or mesenchymal nature (Friedl et al., 2012; Kedrin et al., 2008; Liu et al., 2019; Patsialou et al., 2013) driven by weak intercellular junction strength. The identification of leader versus follower cells in such collective migration and the conceptual and mechanistic overlaps of cancer progression with jamming-unjamming transition has also been an active area of investigation (Sadati et al., 2013).

Multiscale mathematical models that investigate tumor progression have largely focused on angiogenesis and tumor growth. Of late, these models have incorporated cell invasion as a function of cell-cell adhesion and cell-matrix adhesion too (Bearer et al., 2009; Szabo and Merks, 2013). Discrete approaches such as agent- or cell-based ones to study cancer invasion have the advantage of modularizing interactions between the cells and their environment, thus allowing quick transitions from mechanistic hypothesis-framing to rule-based behaviors on cell-population scales (Metzcar et al., 2019). Recent efforts grounded in continuum approaches and lattice growth cellular automaton (LGCA) framework have revealed how heterogeneity in cell-cell adhesion (as a result of many possible reasons: genetic, epigenetic and/or microenvironmental control of EMT or through spatiotemporal variation in cell- and matrix-adhesion dynamics) can pattern the dissemination of cancer cells (Domschke et al., 2014; Reher et al., 2017). Another study combined experiments with simulations to showcase a feedback loop between cell contractility, and the alignment of collagen fibers to posit that intermediate matrix stiffness is optimal for invasion (Ahmadzadeh et al., 2017), thereby emphasizing the nonlinear nature of ECM behavior in determining cancer cell invasion. ECM density and organization, as a function of MMP density, can regulate the switch between proteolytic and non-proteolytic modes of invasion (Kumar et al., 2016). However, the existent literature on cancer invasion, to the best of our knowledge, has not yet explicitly investigated the distinctions and commonalities between mechanisms underlying collective and dispersed modes of invasion. In consequence, phenomena such as mesenchymal streaming (Friedl et al., 2012) and tumor budding (Prall, 2007), which have been meticulously described in pathological and surgical literature, remain largely uninvestigated via mathematical modeling approaches.

Using an experimental setup that mimicked the topographical arrangement of invasive cell clusters encapsulated within BM-like ECM and subsequently by Type 1 Collagen, we were able to observe the cooccurrence of several spatially discernible migratory modes, a phenomenon we called multiscale invasion (simultaneous invasion across multiple scales (Pally et al., 2019)). These experimental observations were simulated using Compucell3D, a modeling framework based on the Cellular Potts model (Das et al., 2017; Glazier and Graner, 1993; Graner and Glazier, 1992; Swat et al., 2012; Zhang et al., 2011), using which we established that the cooperation between a set of biophysical phenomena such as cell-cell and cell-ECM adhesion, cell proliferation and reaction-diffusion-based modulation of matrix proteolysis (computational equivalents of the ECM-degrading matrix metalloproteinases (MMPs) and their inhibitors (The inhibitors of matrix metalloproteinases or TIMPs))-, can give rise to a set of invasive phenotypes that comprise dispersed, collective, and multiscale invasion categories. Here, we investigate how robust each of these phenotypes are. Our multiparametric simulations followed by statistical analyses suggest, to our surprise, that cell invasion is fundamentally of two modes: both are multiscale and subsume exclusively dispersed mesenchymal- and collective-invasions respectively (and hence will be referred to henceforward as dispersed multiscale invasion (DMI) and collective multiscale invasion (CMI) respectively). We explore the inputs that maintain such invasive modes and enable possible interconversions among them, the targeting of which represents insights into novel therapeutic strategies in the future.

## Results

### Construction of an invasion phenospace based on the dynamics of reaction-diffusion, and cell-cell and -matrix adhesion

In our previous manuscript, we were able to demonstrate, using a computational model, that an interplay between parametric combinations of cell-cell and -ECM (BM and Type 1 collagen) adhesion, and reaction-diffusion kinetics of MMP- and TIMP-based ECM remodeling were able to give rise to diverse phenotypes of cell invasion. Not just dispersed- and collective-invasion, but also phenotypes with concurrent presence of the latter two modes were observed, which we termed multiscale invasion. Such simulations were calibrated with *in vitro* 3D culture experiments of breast cancer cells in combinations of appropriate ECMs (Figure 1A). Whereas in our previous effort, proliferation played a dominant role over migration in the invasion of cancer cells, here, we have incorporated active migration due to chemoattraction the fibrillar collagenous ECM (Fang et al., 2014; O’Brien et al., 2010; Postlethwaite et al., 1978; Xu et al., 2019). Our model confirms that invasion can take place in the absence of proliferation (Supplementary Figure 1). We also confirmed that the addition of this ‘active migration’ module did not perturb our previously obtained result that the presence of only BM or only Collagen-like ECM constrained cells to exhibit exclusively collective or dispersed invasion, respectively (Supplementary Figure 2).

**Figure 1:**
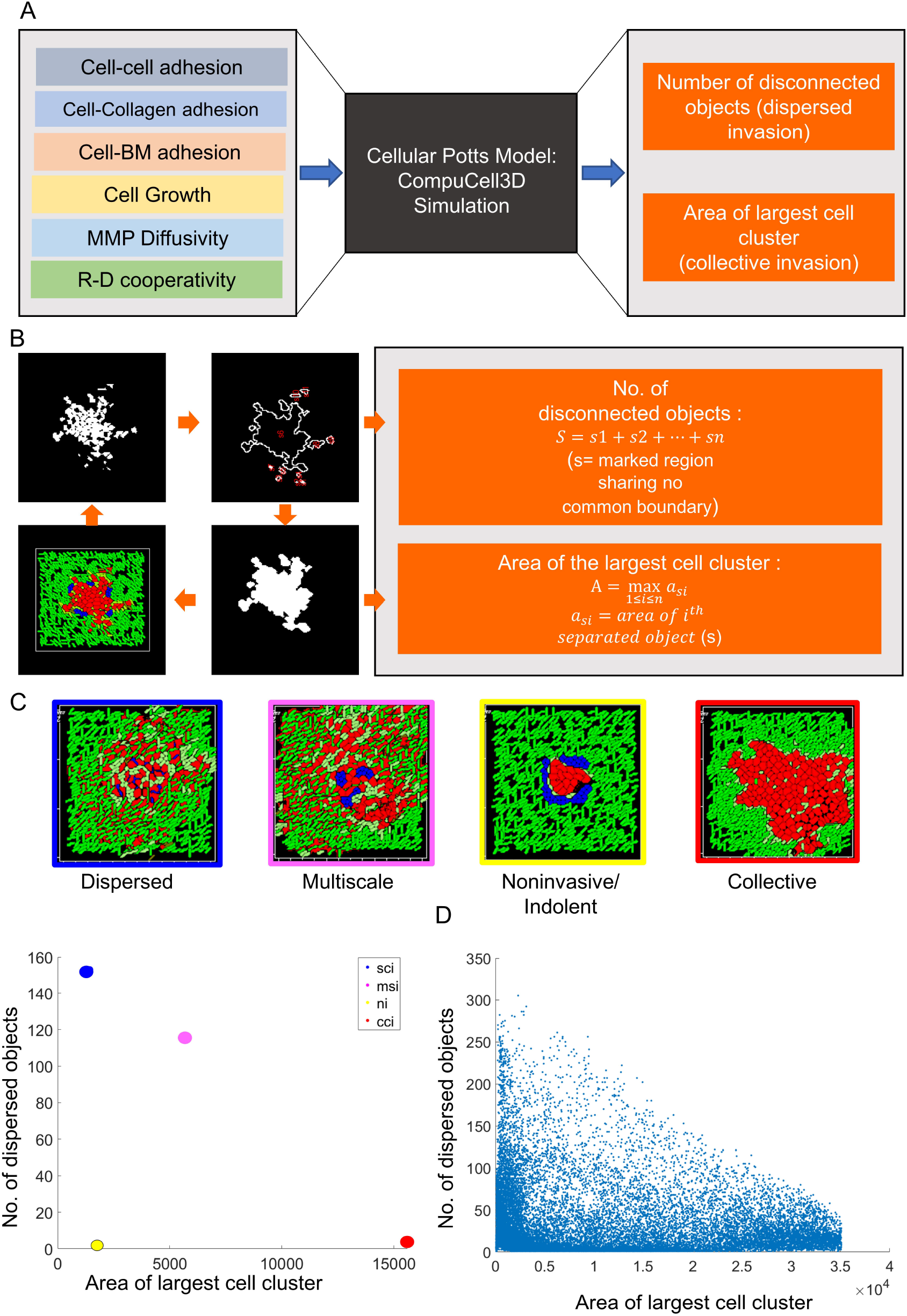
Introduction to the computational framework, simulation inputs and output phenospace. (A) Schematic depiction of the input variables that have been deployed in the CompuCell 3D simulations. The result of the simulations was computed as two outputs that are representative of dispersed invasion and collective invasion. (B) Depiction of computation of the invasion outputs: MATLAB^®^ is used to quantify the morphology of the cancer cell mass in the simulations. 1. Images of all simulations are collected at the 1500^th^ Monte Carlo step (MCS). 2. The images are binarized to isolate the cancer cells. 3. All dispersed ‘objects’ (the objects share no common boundary with each other) are identified and counted and is called ‘No. of dispersed objects’. 4. the mass having largest total covered area is isolated and the value of that highest area is called ‘Area of Largest cell cluster’. (C) Control simulations: Four simulations (red= transformed cancer cell, green= collagen I, blue= Basement membrane (BM)) represent control runs showing morphological variations with certain input variable values. In terms of invasiveness, they are characterized as **dispersed invasion** (blue), **collective invasion** (red), **multiscale invasion** (magenta) and **indolent or non-invasive phenotype** (yellow). The invasiveness of the controls interpreted through the 2 outputs show in 1B allows the construction of a phenospace with X axis measuring the ‘Area of Largest cell cluster’ and Y axis measuring the ‘No. of dispersed objects’. Control simulations can therefore be mapped onto the phenospace of possible output points within the phenospace through various combinations of inputs deployed through simulations run till a similar endpoint (MCS: 1500). (D) Distribution within the phenospace of phenotype outputs as a result of combination of five values for each of the six inputs mentioned in (A) and run three times (replicates). The total number of simulations originating from this combination is 5^6X3=46875. Simulations points having x value more than around 36000 (~90% of the whole simulation lattice) is removed to omit the effect of spatial limitation of the simulation lattice. After that, 99% of the whole dataset (46475 simulations) is retained which is plotted here.

To better quantify the invasion phenotype and characterize the tendency of cells to switch among these phenotypes, here, we constructed an invasion phenospace, wherein the horizontal and vertical axes represent the metrics inherent to collective and dispersed cellular invasion respectively (Figure 1B and C). Therefore, the X axis measures the size of the largest single multicellular mass (the starting ‘primary focus’) at the end of the simulation. The Y axis measures the number of objects (cells or tiny cellular clusters) that are spatially disconnected or dispersed from the primary focal mass at the end of the simulation. Outputs that are closer and parallel to X axis, therefore, represent phenotypes with exclusively collective cell migration; those closer and parallel to Y axis represent dispersed cells migrating solitarily as a result of their disintegration from the primary focal multicellular mass. Outputs with a relatively higher magnitude of values on X and Y axes represent multiscale invasion (MSI) phenotype.

Based on our previous simulation results, we took the following inputs to map the phenospace: a) the contact energies representing cell-cell, cell-BM and cell-collagen adhesion, b) cell division, c) diffusivity of MMP and d) the cooperativity between MMP and its diffusible inhibitor TIMP (the threshold ratio of MMP to TIMP concentration at a given spatial point that is required to degrade Type 1 collagen, which was shown to be an important regulator of multiscale invasion (Diambra et al., 2015; Pally et al., 2019)) (Figure 1A). Whereas this set is, by no means, an exhaustive set of possible inputs, they were shown by previous simulations to significantly impact the phenotypic outcome of invasion and recapitulate the diversity of phenotypes experimentally observed. We chose the input values for which we observed exclusive dispersed, or collective invasion phenotypes and then progressively decreased them in discrete intervals to a minimum limit at which no invasion was observed. In total, 5 values of each input were chosen; outputs were computed for each combination of 5 values of 6 inputs; the computation was repeated 3 times (replicates) using CompuCell3D. The replicates aim to capture the variability in cellular responses enabled by the underlying stochastic framework of the Cellular Potts modeling framework. These outputs were mapped onto the 2D invasion phenospace (Figure 1D). The distribution of outputs indicated that multiscale invasion represented a continuous phenomenon that bridged the exclusive collection of dispersed- and collective invasions close to the respective cartesian axes. Our findings are therefore consistent with conceptual narratives that seek phenomenological continuity between invasive modes but suggest that the transition between such modes can occur through a phase plane rather than in a linear manner. Whereas our output method does not enable the identification of optimal parametric combinations based on design space hypercube (as done by (Ozik et al., 2019)), it allowed us to examine simultaneously two model outputs simultaneously which could describe the predominant cell invasion phenotypes.

### Identification of optimal clustering of the invasive phenospace

We next investigated the optimal number of phenotypic clusters that can best classify the scatter of outputs in the two-dimensional phenospace. Such approaches are commonly used for gene expression studies to unravel hidden patterns in multivariable high-dimensional spaces (Oyelade et al., 2016). K-means algorithm is a commonly used unsupervised clustering method which, given *a priori* an integer value of K, partitions the given dataset into K disjoint subsets (Macqueen, 1967). It selects a random initial seed point of preferred clusters which are used to cluster the remaining data points; thus, different runs of K-means can possibly give rise to distinct clusters and their arrangements.

We performed K-means clustering on the phenospace for K=2 to K=8 for multiple instances, and as expected, observed different clusters across multiple (n=15) runs for a fixed value of K, except for K=3 (Figure 2A and Supplementary Figure 3). For K=2, two predominant categorization patterns were seen. In the first pattern, which was obtained slightly more than 50%, one of the clusters indicated phenotypes with relatively lower invasion (blue outputs in Fig 2A, top row, left panel, #1); the other cluster consisted of points with moderate to high values of both dispersed and collective invasion (yellow clusters in Fig 2A, top row, left panel, #1). In the other categorization pattern for K=2, we observed a cluster denoting largely collective cell migration (blue cluster in Fig 2A, top row, right panel, #2), and another one encompassing the rest of the outputs (yellow cluster in Fig 2A, right panel, #2). For K=3, the phenospace clustered into what could be characterized as non-invasive to indolently invasive phenotypes (blue cluster in Fig 2A, middle row panel), and two invasive phenotypes (yellow and green cluster in Fig 2A), each of which included, but were not limited to, purer single cell- and collective invasion-representing outputs close and parallel to the Y and X axes respectively and will, therefore, be referred to as dispersed multiscale invasion (DMI) and collective multiscale invasion (CMI) respectively. This pattern was obtained 15 out of 15 times for K=3 clustering, suggesting the robustness of this categorization for further analysis. For values of K>3, we observed two or more cluster patterns for independent runs of K-means clustering (Figure 2 and Supplementary figure 3A). Clusters within such patterns could be subsumed within the two distinct multiscale invasion clusters seen for K=3 (see for e.g., cluster pattern #2 of K=4), or in other cases spatially independent of the latter (for e.g., cluster pattern #1 of K=4).

**Figure 2:**
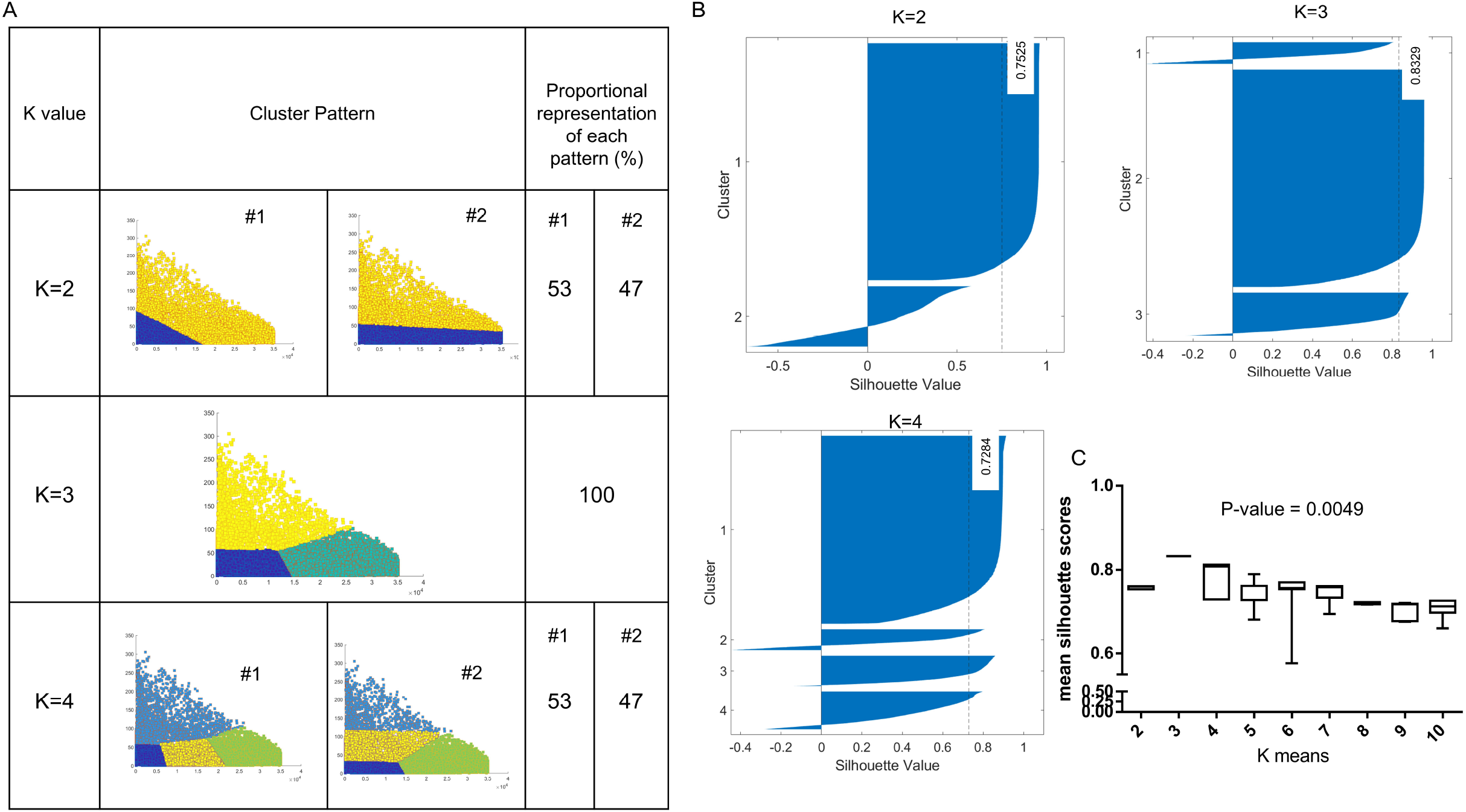
The invasion phenotypic outputs can be optimally segregated into three clusters. (A) All the simulation outputs in the phenospace (in Figure 1D) are analyzed with K-means clustering using cluster number K=2-4 (leftmost column). Clustering was performed 15 times with random initial cluster centers. The cluster phenotype patterns are shown (middle column) along with their proportional representation within the 15 replicates (right column). (Cluster patterns for K=5-8 shown in supplementary figure 3) (B) The optimal value for K computed using the Silhouette method. Silhouette plots for K=2-4 are constructed with X axis signifying silhouette values and Y axis representing cluster numbers. (Silhouette plots for K=5-8 shown in supplementary figure 3) (C) The average silhouette values and associated standard deviation for all 15 clustering replicates in different K-value groups are plotted in the box and whisker plot (bottom right). Statistical test was performed using unpaired ANOVA with Tukey’s post hoc multiple comparison (p=0.0049).

Next, we conducted Silhouette analysis on K-means to quantify the consistency in clustering, for varied values of K. Silhouette scores are indications of how similar a data-point is to others in its own clusters, compared to those in other clusters. Thus, Silhouette scores provide a measure of both the tightness of the clusters defined as well as the extent of separation among them. These values range from −1 to +1; the higher the value, the closer a given datapoint is to other points in those clusters and the farther it is from other clusters (Zhao et al., 2018). Thus, a higher average score would indicate a greater degree of cluster separation. Clustering for K=3 reported a higher average Silhouette score as compared to K=2, 4, 5, 6, 7 and 8 (Fig 2B and Supplementary Figure 3B). Moreover, over 15 replicates of K-means tried over the same phenospace, the mean and variance of the average Silhouette scores was calculated. K=3 showed the highest mean and the least variance (Figure 2B), further endorsing that K=3 suggests the optimal clustering scenario for the phenospace. We asked if the outputs within what we labelled as the noninvasive or indolent cluster indeed showed lower dissemination relative to the two multiscale modes. To answer this, we mapped the area of the smallest circle that could enclose all the cells for a given output within the phenospace. Comparing the mean invasion revealed that CMI cluster showed the greatest spatial spread followed by DMI; the indolent cluster indeed invaded very poorly with respect to the other two (Supplementary Figure 4)

### The two multiscale invasion clusters exhibit distinct phenotypic patterns

Having identified the two multiscale invasion clusters, we asked how morphological phenotypes changed between these clusters. To do so, we chose representative outputs within each cluster (in the middle of a cluster, near its boundary with the other cluster or close to the X/Y axis) and examined the endpoint of simulation (1500^th^ Monte Carlo Step (MCS) of the simulations). The output within the indolent cluster (close to X=0, Y=0) showed scarce migration of cells within the fibrillar ECM (Figure 3A). For representative outputs close to the boundaries between indolent cluster and the CMI and DMI clusters, the invasion phenotypes represented fingerlike cellular streams with dispersed single cells around them reminiscent of fingering instabilities seen for immiscible fluids of distinct densities (Aref and Tryggvason, 1989; Mikaelian, 1990) (Figure 3B and C).

**Figure 3:**
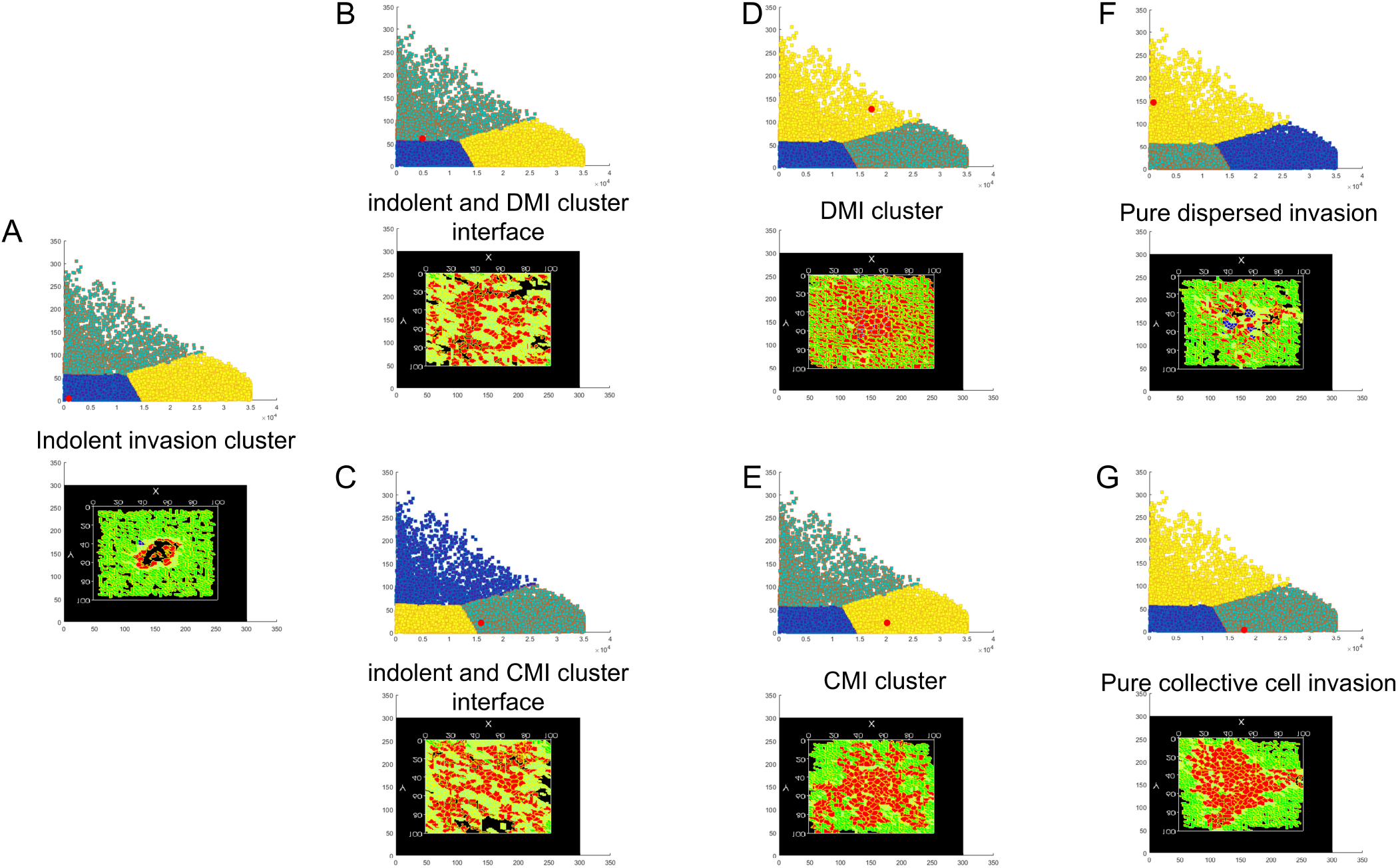
Visualization of distinct invasive phenotypes across the phenospace. The simulation images on the bottom part of each subfigure is at MCS 1500 and their respective position is denoted with red color in the (K=3)-means clustered phenospace above. (A) The end point of a simulation output chosen within the indolent cluster close to the X=0, Y=0 shows scarce invasion of cells from the originating locus. (B-C) The end point of simulation outputs chosen at the indolent-DMI cluster boundary and the indolent-CMI cluster boundary respectively, shows connected networks of cells with oligocellular dispersed masses, reminiscent of tumor budding (Prall, 2007). (D) The end point of a simulation output chosen inside the DMI cluster shows a phenotype, wherein a growing central mass of cells is surrounded by a plethora of dispersed cells. (E) The end point of a simulation output chosen inside the CMI cluster shows a phenotype, wherein the growing central mass invades through finger like projections with a few dispersed cells close to these projections. (F) The end point of a simulation output chosen inside the DMI cluster close to Y axis shows a classical dispersed invasive phenotype. (G) The end point of a simulation output chosen inside the CMI cluster shows a classical collective cell invasive phenotype.

A representative output in the center of DMI cluster showed a centrally located mass invading collectively surrounded by dispersed cells in the fibrillar collagenous ECM (Figure 3D). On the other hand, in the center of the CMI cluster, the representative output showed connected streams of cells interspersed with few single invading cells in fibrillar ECM (Figure 3E). For the output in the DMI cluster close to the Y axis, all the cells that had started out as a collective, were dispersed into the fibrillar ECM (Figure 3F). Its counterpart in the CMI cluster, close to the X axis, showed a dysmorphic bulk of cells growing centrifugally in a collective manner into the fibrillar ECM (Figure 3G). In summary, the DMI and CMI clusters represent multiscale invasions with a greater degree of cell dispersal and intercellular networking, respectively. Our study concurs with previous experimental examinations of cancer cell phenotypes in 3D, wherein cell lines within BM matrices exhibited mass-, grape- or stellate morphologies (Kenny et al., 2007). However, the addition of a collagen-like fibrillar ECM in our computational model unmasks stunning phenotypic diversity within the stellate morphology through an operational dialectic between cellular connectedness and dispersal.

### Determinants of the different phenotypes observed in invasive phenospace

We next asked which of the inputs (Figure 1A) may be proportionately greater represented within each of the three clusters. In order to do so, we generated inputoutput maps of our phenospace, wherein each output was color-coded based on the value of input that contributed to it (highest two input values were denoted in red color, intermediate value blue, and the lowest two values were denoted with green color). We observed that the lowest and highest values for cell-cell adhesion contact energies were maximally and minimally apportioned within the indolent cluster respectively. This implied that input combinations comprising high cell-cell adhesion contributed strongly to the indolent phenotype (Figure 4A; Table 1). This is consonant with the demonstration that strong adhesion mediated through intercellular junctions contributes to cellular and tissue architecture (Bhat and Bissell, 2014). On the other hand, high values of cell-collagen and cell-BM adhesion significantly contributed to DMI and CMI cluster outputs respectively (Figure 4B-C, Table 1). This is consonant with experimental demonstrations of the necessity of cancer cells to adhere to ECM substrata for migration (Gkretsi and Stylianopoulos, 2018). Lowest values of both inputs were proportionately seen to be higher within the indolent cluster output phenotypes. We have also observed computationally that the varying contributions of BM or Collagen around cancer cells can potentiate collective and dispersed invasion respectively (Supplementary Figure 2; the input signatures across these two clusters seem to be distinct as well, Supplementary Figure 5). Higher values of cell proliferation contributed strongly to CMI phenotype but in comparison, were depleted for DMI cluster outputs (Figure 4D, Table 1). In comparison, parameters cognate to reaction-diffusion kinetics: diffusion rate of MMP and cooperativity between MMP and its inhibitors in MMP degradation showed mild variation in their tendency to be apportioned between the three clusters: highest values of MMP diffusion rates were represented relatively to the greatest extent within DMI cluster, whereas both multiscale invasion clusters were characterized with a relative enrichment of outputs with low MMP/TIMP cooperativity values (Figure 4E-F; Table 1). To summarize, the DMI cluster was composed of outputs associated with high cell proliferation and cell-BM adhesion. The CMI cluster was associated with high cell-Collagen adhesion, whereas indolent invasion cluster consisted of outputs with high cell-cell adhesion

**Figure 4:**
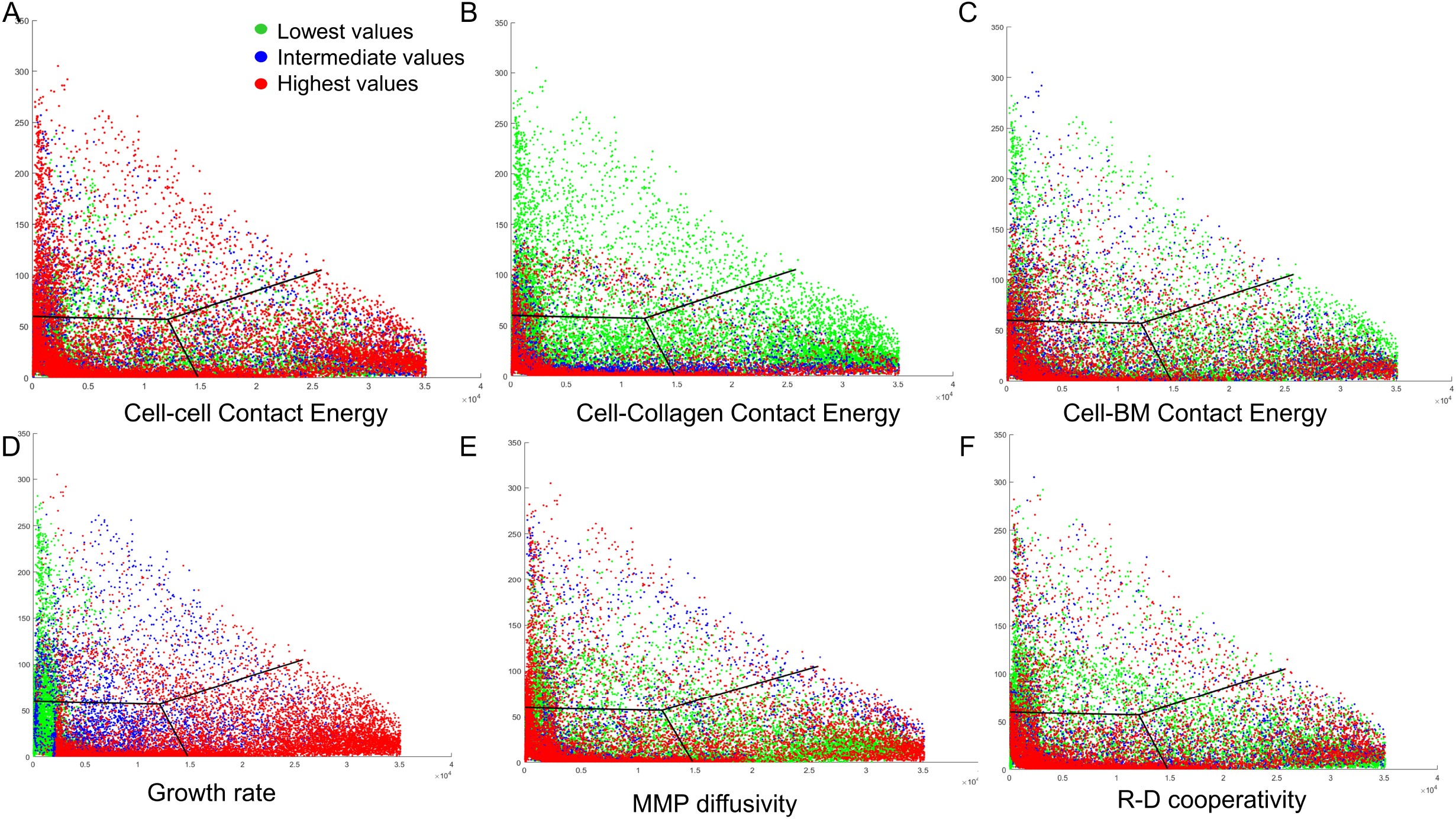
Coarse-grained classification of simulations in the phenospace based on input variable values: The phenospace is divided into 3 parts depending on the K=3-means clustering and the boundary between the clusters is denoted by the black lines. The input variable values (for cell-cell contact energy (A), cell-collagen contact energy (B), cell-BM contact energy (C), growth rate (D), MMP diffusivity (E), and R-D cooperativity (F) (see also Materials and Methods; parameter scan section) attributed to the simulation points in the phenospace are collected and after analysis, all the phenospace points are colored based on the values of the respective input variables. The lower two values are denoted in green color, the intermediate value in blue, and the higher two values are denoted in red color.

**Table 1:**
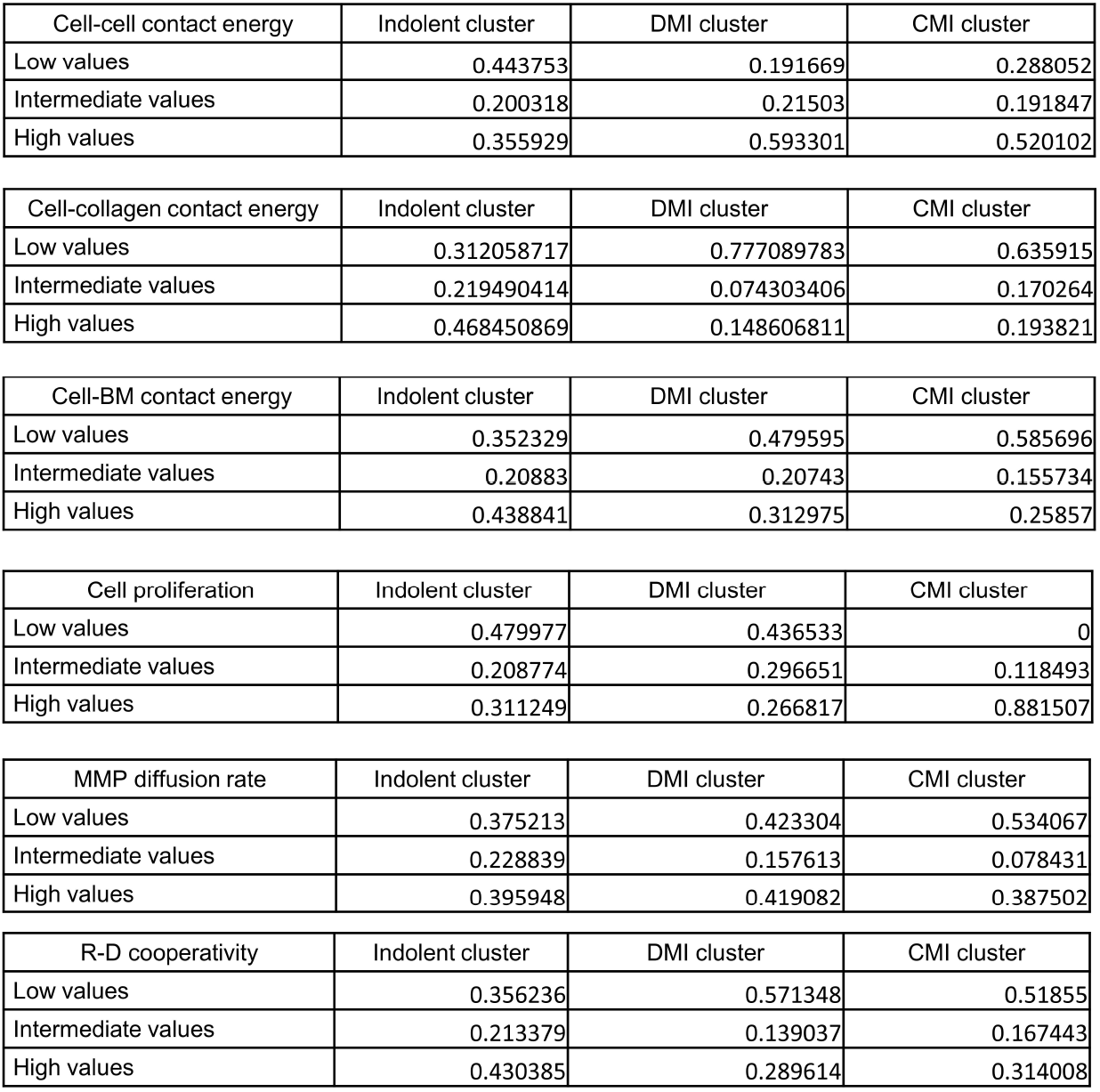
Cluster-wise fraction of input variable values. Table showing the proportion of two lowest input values, intermediate value and the two highest input values for six inputs within the three clusters representing indolent invasion, collective multiscale invasion (CMI) and dispersed multiscale invasion (DMI).

| Cell-cell contact energy | Indolent cluster | DMI cluster | CMI cluster |
| --- | --- | --- | --- |
| Low values | 0.443753 | 0.191669 | 0.288052 |
| Intermediate values | 0.200318 | 0.21503 | 0.191847 |
| High values | 0.355929 | 0.593301 | 0.520102 |

| Cell-collagen contact energy | Indolent cluster | DMI cluster | CMI cluster |
| --- | --- | --- | --- |
| Low values | 0.312058717 | 0.777089783 | 0.635915 |
| Intermediate values | 0.219490414 | 0.074303406 | 0.170264 |
| High values | 0.468450869 | 0.148606811 | 0.193821 |

| Cell-BM contact energy | Indolent cluster | DMI cluster | CMI cluster |
| --- | --- | --- | --- |
| Low values | 0.352329 | 0.479595 | 0.585696 |
| Intermediate values | 0.20883 | 0.20743 | 0.155734 |
| High values | 0.438841 | 0.312975 | 0.25857 |

| Cell proliferation | Indolent cluster | DMI cluster | CMI cluster |
| --- | --- | --- | --- |
| Low values | 0.479977 | 0.436533 | 0 |
| Intermediate values | 0.208774 | 0.296651 | 0.118493 |
| High values | 0.311249 | 0.266817 | 0.881507 |

| MMP diffusion rate | Indolent cluster | DMI cluster | CMI cluster |
| --- | --- | --- | --- |
| Low values | 0.375213 | 0.423304 | 0.534067 |
| Intermediate values | 0.228839 | 0.157613 | 0.078431 |
| High values | 0.395948 | 0.419082 | 0.387502 |

| R-D cooperativity | Indolent cluster | DMI cluster | CMI cluster |
| --- | --- | --- | --- |
| Low values | 0.356236 | 0.571348 | 0.51855 |
| Intermediate values | 0.213379 | 0.139037 | 0.167443 |
| High values | 0.430385 | 0.289614 | 0.314008 |

### Perturbing input-output combinations to identify transitions in the phenospace

We sought to test the strength of these associations by perturbing these specific input values within these clusters. We minimized the cell-BM adhesion (through increase in cell-BM contact energy) in 100% of outputs in the CMI cluster that showed highest values of this parameter. Whereas 38% outputs remained within the CMI cluster after decreasing adhesion, 54% outputs transitioned to the indolent cluster and 8% to the CMI cluster (Figure 5A). Minimizing growth rate in outputs from DMI associated with its highest values brought about a transition of 96% outputs to the indolent cluster and only 4% to the CMI cluster (Figure 5B). Decreasing cell-collagen adhesion in 100% of outputs in the DMI cluster that showed highest values of this parameter caused 88% outputs to transition to indolent cluster with only 2% transitioning to the CMI cluster (Figure 5C). Interestingly, maximizing values of R-D cooperativity for outputs showing the lowest values within the CMI cluster brought about a transition in 18% outputs to the DMI clusters (Figure 5D). Similarly, R-D cooperativity and MMP diffusivity were increased and decreased in DMI cluster outputs that showed the lowest and highest values of these two input traits, respectively; transitions of 17% and 34% were seen as a consequence of such perturbations. This indicated that parameters that render the two multiscale invasive phenotypes unique, the matrix adhesion parameters when perturbed, lead to a loss of invasion. On the other hand, inputs relating to the reaction-diffusion-based proteolysis of ECM had a greater probability of allowing transitioning between of cells between CMI and DMI phenotypes. These representative examples identify how cells can traverse the phenospace through these specific alterations in the input parameter(s).

**Figure 5:**
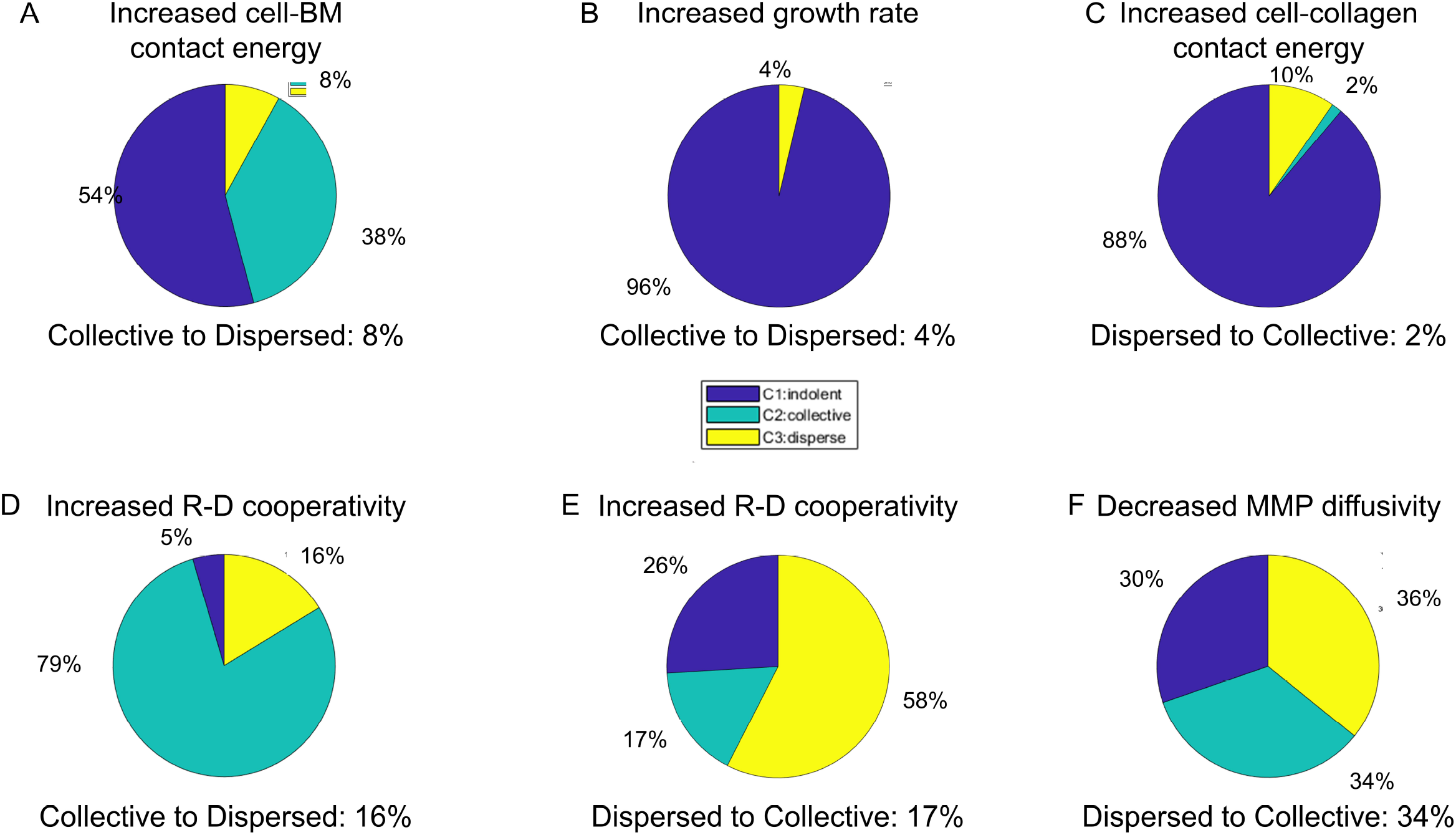
Transition of points between phenotypic clusters is controlled by the input variables. (A) Pie chart showing the transition of outputs from the CMI cluster showing highest cell-BM adhesion, upon decreasing the latter to a minimum value. (B) Pie chart showing the transition of outputs from the CMI cluster showing highest growth rates, upon decreasing the latter to a minimum value. (C) Pie chart showing the transition of outputs from the DMI cluster showing highest cell-collagen adhesion, upon decreasing the latter to a minimum value. (D) Pie chart showing the transition of outputs from the CMI cluster showing lowest R-D cooperativity values, upon increasing the latter to highest values used in the paper. (E) Pie chart showing the transition of outputs from the DMI cluster showing lowest R-D cooperativity values, upon increasing the latter to highest values used in the paper. (F) Pie chart showing the transition of outputs from the DMI cluster showing highest MMP diffusivity values, upon decreasing the latter to a minimum value (Proportion of outputs in the pie charts within the indolent, DMI and CMI cluster shown in blue, yellow and green).

## Discussion

In an illustrative review, Friedl and coworkers provided an elegant classification of generic cell invasion: they enumerated numerous movements with solitary and bulk movements at the two ends of the phenotypic spectrum (Friedl et al., 2012). In addition, a three-dimensional graph shows how the three axes of leading-edge polarity, apicobasal polarity and cell-cell contacts can accommodate the different modes of phenotypes. This idea is based on an earlier review on the same topic where Friedl and Wolf argue for a “multiparameter tuning model of invasion wherein a combination of cues such as density, stiffness, and orientation of the extracellular matrix together with cell determinants—including cell–cell and cell–matrix adhesion, cytoskeletal polarity and stiffness, and pericellular proteolysis—interdependently control migration mode and efficiency” (Friedl and Wolf, 2008; Wolf et al., 2007). Our manuscript seeks to build on this idea by subjecting it to mathematical rigor and identify its underlying assumptions and mechanisms. While Friedl and others have acknowledged that distinct modes may concur spatially, their tuning model would attribute such co-occurrence to the heterogeneity in microenvironmental cues. However, does the co-occurrence of dispersed and collective modes of invasion (multiscale invasion or MSI) represent a distinct invasive mode, wrought through distinct signatures from a tunable combination of inputs? In this manuscript, we argue that such is the case. In fact, the input combinations give rise to not one but two types of MSI. The two types have higher relative representation of dispersed and collective cell invasion respectively and explain the exclusively solitary and bulk invasion respectively.

Invasion of cancer cells has been characterized as non-equilibrium phase separation of mesoscale multicellular aggregates (Yang et al., 2019). This idea is an extension of previous frameworks which describe spreading cellular ensembles as liquids and dispersed cells as 2D gases (Sadati et al., 2013; Trepat and Fredberg, 2011). Jammed non-invasive states have been likened to ‘solid phases’ and the transitions between non-invasive and invasive (collective and dispersed) phenotypes can be compared with phase transition dynamics. The novelty of our contribution within this framework stems from the crucial addition of two ECM microenvironments instead of one as well as the recreation of the collagenous ECM by invading cancer cells. This is not to add needlessly to the complexity but to incorporate essential characteristics of tumorigenic epithelial tissues (Naba et al., 2014; Nelson and Bissell, 2005). The production of fresh collagenous ECM is being increasingly demonstrated to play a crucial role in defining the invasive behavior of cancers. The ECM secreted by cancers is rich in fibrillar collagen. In a recent preprint, we identify unique physicochemical features associated with the collagenous ECM secreted by invasive breast cancer cells (https://doi.org/10.26434/chemrxiv.12063420.v1). Such observations have been corroborated in several cancers including that of breast (Naba et al., 2014). Upon incorporation of these two features, we observe that the phenotypic distribution that is intermediate between pure collective and dispersed states can be fundamentally split into two clusters, with specific input contributions. Our findings allow us to extend the characterization of mesoscale multicellular phenotypes based on soft matter models (Gonzalez-Rodriguez et al., 2012), as possibly polydisperse media with DMI comparable to aerosol and CMI comparable with foam states, respectively.

Surgical histopathological literature is replete with observations of multiscale behavior in cancer invasion. In the context of colorectal carcinoma, Prall and coworkers have made meticulous observations on ‘tumor budding’, wherein single cells or small collectives of, cells seem to escape from growing edges of malignant neoplasms (Prall et al., 2005; Prall et al., 2009). Such multiscale behavior is associated with a more sinister kinetics of metastasis and poor prognosis (Hase et al., 1993; Prall et al., 2005; Prall and Ostwald, 2007). Tumor budding has been observed in breast and pancreatic cancers as well (Petrova et al., 2019; Salhia et al., 2015) and have been connected to a partial EMT phenotype (Bronsert et al., 2014; Grigore et al., 2016). Our previously published experiments, which were crucial to the development of the computational model we have explored here, described multiscale invasion, which is phenomenologically similar to tumor budding. Indeed, early investigations by others confirm our empirical and theoretical findings that surface proteins and ECM play key roles contributing to multiscale invasion (Graves et al., 2016; Masaki et al., 2003). In our present analysis as well, we observe the phenotypic subspace pertaining to truly multiscale invasive phenotypes show the highest proportion of dissemination, compared with subspaces associated with pure collective and dispersed invasion (Supplementary Figure 4).

At this point of time, our study does not incorporate three salient aspects of invading cancer cells. The first is the change in shape of cells as they move through matrix microenvironments. This distinguishes amoeboid from mesenchymal single cell migration and is associated with shape-based asymmetries in remodeling of ECM intercellular adhesion (Pankova et al., 2010). Despite this lacuna, we believe our dispersed multiscale invasion is mesenchymal in nature as it is predicated on adhesion of cells to the ECM (which distinguishes it from amoeboid migration), and appropriate for the level of graining that is achieved in our analysis. However, cell shape dynamics will be implemented in our future efforts. Secondly, our computational environment does not incorporate the stromal cells that contribute in significant ways to the phenomenology of cancer cell invasion (Labernadie et al., 2017) due to limitations of the sheer computational power required to compute the dynamics of a multi-cell multi-ECM environment. We assume that the R-D dynamics of MMP-TIMP incorporated in the computational environment is a downstream effect of the cancer and stromal cell activities. This assumption will be tested in future efforts. Last, but not the least, our model does not incorporate cell polarity which has been proposed to contribute to active models of soft biological matter (Perez-Gonzalez et al., 2019) and predict migration in the absence of proliferation (Tlili et al., 2018). Future efforts will be devoted to computational investigating the properties of polarized polydisperse states. At present though, our effort serves to put multiscale migratory behavior in the fundamental center of the concept of cancer invasion. Understanding the process has potentially far reaching consequences for future therapeutic efforts.

We would conclude by drawing a parallel between our study and the modeling efforts employed for investigating developmental processes (Alber et al., 2006; Alber et al., 2004; Christley et al., 2007; Harrison et al., 2011; Izaguirre et al., 2004; Kiskowski et al., 2004; Merks and Glazier, 2006). While our model derives its motivation, initial conditions and tissue geometry (which have been shown to be a crucial determinant in morphogenetic pattern formation (Nelson et al., 2006)) from breast cancerous contexts (Bhat and Bissell, 2014), one can as well interpret our model as a developmental system wherein inter-agent adhesion and reaction-diffusion based phenomena sculpt cellular patterns. In fact several of our model inputs represent the biological physics associated with the framework of dynamical patterning modules, which has been employed to study the evolution and logic of developmental mechanisms (Newman and Bhat, 2008, 2009). In effect, this suggests that the redeployment of developmental and morphogenetic principles in the context of genomic aberrations may underlie the mechanisms behind discrete invasive behaviors of cancer cells and the transitions between them.

## Materials and methods

### Modelling framework

Compucell3D (CC3D) is a problem-solving environment that combines the lattice-based GGH (Glazier–Graner–Hogeweg) model or CPM (Cellular Potts model) with PDE solvers and other models to allow for study and simulation of multiscale virtual biological processes (Swat et al., 2012). The software divides the whole simulation lattice into ‘cells’ (collection of pixels). A specific ‘cell type’ is assigned to each of them. Interaction parameters between cell types can be made to approximate biological constraints between components, similar to that of the original in vitro or in vivo biological system. Such constraints or parameters regulate the simulation through the effective energy or **Hamiltonian (H)** calculated at each **Monte Carlo Step (MCS)**. Calculation of H determines the allowed configuration and behavior of cell*s* at each MCS.

Hamiltonian or effective energy calculation at each MCS:

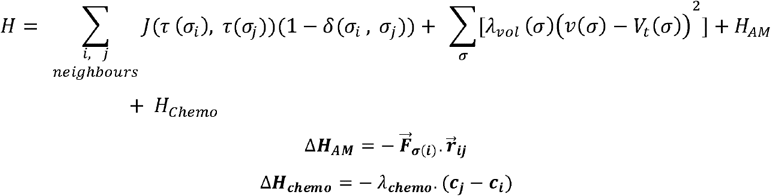

Index(σ)-copy attempt (from pixel i to j) success or rejection incorporating Boltzman probability function:

For Δ***H***≤ 0 condition, the associated index-copy attempt will be successful, so the target pixels are updated. So, success probability is *P* = 1

For Δ***H***> 0 condition, the associated index-copy attempt will be successful with a probability of 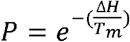 and it will be unsuccessful with a probability *P*’ = 1 – *P*

There are 4 contributors for calculating the H for cells.

The first contributor is the sum over all neighboring pairs of lattice sites *i* and *j* with associated contact energies (*J*) between the pair of cells indexed at those *i* and *j*. In this term, *i, j* denotes index of pixel, σ denotes cell index or ID, and *τ* denotes cell-type. The δ function in this term will ensure that only the σ_*i*_ ≠ σ_*j*_ terms are calculated (i,j belonging to same cell will not be considered). Contact energies are symmetric [*J* (*τ*(*σ_i_, τ*(*σ_j_* = *J*(*τ*(*σ_j_*),*τ*(*σ_i_*))]. The contact energy between two cell*s* is considered to be inversely proportional to adhesion between those two cell*s*.

The second contributor is a function of the volume constraint on the cell, where for the cell σ, λ_vol_(σ) denotes the *inverse compressibility* of the cell, v(σ) is the number of pixels in the cell (*volume*), and *V*_t_(σ) is the cell’s *target volume.* For each cell, this term is governed by its growth equation.

The third term related to random active motility. For calculating Δ***H**_AM_* for a given MCS step, when an index-copy attempt is made for pixels from “i” to “j” for a cell *σ*(*i*), the force vector is 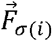 and the distance vector between those pixels is 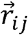. Therefore, the product of these two vectors with correct direction alignment will satisfy the condition to minimize H and will determine the movement of the cell along that direction. Effectively, the respective steppable (CellMotilitySteppable) provides an external force on the centre of mass of the cell*s* which changes direction randomly every MCS.

The fourth term is relevant for biased motility of the cell due to chemotaxis. For calculation of Δ***H***_*chemo*_, *c_j_* represents the concentration of the chemical field (‘GF’ for our simulations) at index-copy target pixel (j) and *c_i_* represents the concentration at index-copy source pixel (i). *λ_chemo_* is a constant which determines how strongly the cell will respond to the external chemical gradient by its value and its sign determines the whether the cell should move towards positive or negative gradient of concentration.

In the Boltzman probability function, *ΔH* represents the calculated change in overall Hamiltonian of the system between the system-configuration at previous MCS and a specific system-configuration at current MCS. T_m_ relates to effective membrane fluctuation for the cell. (https://compucell3dreferencemanual.readthedocs.io/en/latest/index.html) (Swat et al., 2012).

#### Simulation lattice

We use a 100* 100*1-pixel square lattice with non-periodic boundary for all the simulations (the initial configuration of simulations resembles Fig 1(C) where no invasion of any type is observed).

#### Cell types

There are total 6 different cell types used in the simulations-

##### Medium

all cells with unassigned cell type are medium cells. These cells act as free, uninterrupted space in the simulation space.

##### Cancer

these cell*s* are initially situated at the center grid surrounded by basement membrane (BM). Cancer cells start as rectangular objects of 16-unit volume (4 × 4 pixels), spanning a total 14 × 14 × 1 unit volume (16 cancer cell*s*) at the center (*x, y* = 43:57) of the simulation grid without any intercellular space. Cancer is the only cell type that is allowed to grow and proliferate. To differentiate from non-cellular components, cancer cell*s* have membrane fluctuation unlike others. The cell*s* also have random motility associated with them. An ‘ExternalPotential’ plugin was used to enable ‘cellMotility’ to act on the cancer cell*s* through the Hamiltonian.

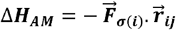

for a given MCS step, when an attempt is made to copy a pixel from “i” to “j” for a cell *σ*(*i*), the force vector is 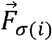 and the distance vector between those pixels is 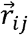. Therefore, the product of these two vectors with correct direction alignment will satisfy the condition to minimize H and will determine the movement of the cell along that direction. The steppable provides an external force on the center of mass of the cell*s* which changes direction randomly every MCS, therefore mobilizing the cell*s* in different directions and providing them with active motility like biological cells. The direction of 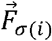 is randomly changed at any angle for each cell in each MCS; thereby favoring motility in random directions for all cell*s* for a short period and changing direction again. In the ‘cellMotilitySteppable’ of cancol2Steppable.py file, in ‘def start(self)’ we have randomly initialised the force vectors.

##### BM (laminin)

This cell type surrounds the cancer cell*s* in the initial configuration. 2 layers of tightly packed BM cell*s* (x, y = 37:63; 3 × 3 pixels= 9 unit volume) separate the cancer cell*s* from C1 cell*s*. To approximate extracellular matrix (ECM) architecture surrounding luminal epithelial cells in mammary duct, BM cell*s* are modelled as dense adhesive bloblike objects to mimic the lamina densa of basal lamina.

##### (Collagen) C1

Outside BM cell*s*, fibrillar C1 cell*s* mimics the interconnected fibrils of collagen I. C1 cell*s* span the remaining space of the simulation lattice (x, y = 63:100). Interfibrillar gaps are characteristic of the C1 cell region unlike BM. The gaps aid in non-proteolytic cancer cell motility through C1 layer. In the initial configuration of the simulation, elongated C1 cell*s* are orientated in random directions with each cell*s* having 4 x 2 =8 unit volume.

##### C_lysed

This cell type is used in an intermediate step during matrix degradation and regeneration. Reaction-diffusion dynamics of the chemicals secreted by cancer cell*s* allows for degradation of BM and C1 matrix cell*s*. Upon meeting certain criteria for degradation, the BM or C1 cell type of a particular cell becomes C_lysed, although retaining the shape and size of that cell. These cell*s* track the MCS from their individual degradation event and transform into newly synthesized matrix cell*s* after 20 MCS. They have properties specified as intermediates of C1 and medium cell types. C-lysed type signifies the degraded ECM, predominantly Type 1 Collagen fibers. Collagen fibers upon degradation are known to produce substrates that act as attractant cues for chemotaxis (Postlethwaite et al., 1978). In addition, collagenous matrices have been shown to be depots of growth factors (Schuppan et al., 1998; Somasundaram et al., 2002; Somasundaram et al., 2000).

##### NC1

Designed to mimic the ‘cancer matrisome’, the newly synthesized matrix cell*s* are denoted as NC1. These cells are almost like C1 in their behavior and can undergo further degradation to become C_lysed and subsequently after 10 MCS would become NC1 again. If undisturbed the C_lysed and NC1 cell type transformations keep the position, size and shape of the cell unchanged.

The exact difference between C1 (stromal collagen) and NC1 (newly synthesized collagen) are as follows:

1. There is a volume reduction we have implemented in NC1, to account for diffusion of soluble degraded ECM, 0.005 unit vol./MCS * 20 MCS= 0.1unit
2. The chemoattractant GF is secreted by C1, when it comes in contact with C_Lysed i.e. during degradation. But even though NC1 can be degraded, it does not secrete GF. Both of them gets converted into C_Lysed and that cell-type secretes GF.
3. Among both, only the contact energy term between cell and C1 is an input variable for parameter scan.

#### Contact energies

As the whole simulation lattice is divided into cell*s*, the contact energy is used to regulate the sorting of all the cell*s*. Specific contact energy values are assigned to all pairs of interactions between different and same cell types. The contact energy is also *inversely proportional* to adhesion between the respective cell type pair or components of the system. As there are a total of 6 cell types in the simulation, there are total 21 contact energies that need to be assigned and the values determine the differential adhesion. The values of the contact energies were set using control and validation experiments from our previous publication (Pally et al, Frontiers, 2019). 3 among those 21 contact energy- or adhesion-parameters were selected as input variables, which would be changed during further analysis. These 3 contact energies or adhesions are associated with pairs of cell types of only cancer cell*s* (**c-c CE**), cancer cell*s* and BM (**c-lam CE**), cancer cell*s* and C1 (**c-c1 CE**) since in our previous efforts we had identified the combination of other contact energies (such as those between BM and Type 1 collagen that were permissive for invasion-based simulations.

The neighbourhood (‘NeighborOrde?) for adhesion term is 2. This ensures for a square shaped cell, all neighbouring cells on its 4 sides along with cells on the 4 corners are considered during the Hamiltonian calculation.

#### Reaction diffusion

CC3D allows chemical fields to determine specific spatiotemporal *cellular* behaviour during simulation. The fields contain the values of concentration of the chemical at each location of the simulation grid. Two chemicals, A and I, are used as activator and its inhibitor as per reaction diffusion dynamics and their concentrations are governed by the partial differential equations (PDE). The governing equations for these two fields are:

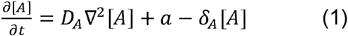

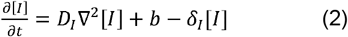

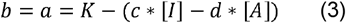

Where, [A], [I]: concentration values for fields A and I.

*D_A_,D_t_*: diffusion constants of A and I

*δ_A_, δ_I_*: degradation rates of A and I

a, b: secretion rate of A and I

t ≡ MCS

Default parameterizations, *D_A_*= input variable, *D_I_*= 0.04, *δ_A_*= *δ_I_*= 0.003, K= 2.0, c= 4.0, d= 2.0

Here A is considered as the activated form of matrix metalloproteinases (MMPs) and I is considered as tissue inhibitor of matrix metalloproteinases (TIMP). The difference in diffusion constants of A and I formulates the pattern in Turing space, so altering one of them (in this case, *D_A_*) can result in different RD dynamics for different simulations. Activation (or secretion of the activated form i.e. ‘a’) of A is assumed to be dependent on its inhibitors (inversely) and on its own concentration (autocatalysis). The cancer cell*s* secrete both A and I when they come in contact with matrix cells such as C1, BM and NC1. Their concentrations are also calculated at the center of mass of the matrix cells.

If the ratio ([A]/[I]) of concentrations of A and I at the center of mass of any matrix cell is more than a threshold, then that matrix cell is degraded. The threshold value for the ratio is termed as RD cooperativity and is one of the input variables. The value of RD cooperativity signifies the inhibition effect of the inhibitor on the activator’s activity (matrix degradation) during the course of a simulation. After degradation and conversion to C_lysed cell type, the cell becomes NC1 type in 10 MCS which undergoes same treatment as matrix cell*s* again. This regeneration of matrix is essential to eliminate unnecessary free spaces formed as an artefact of matrix degradation which takes the computational model closer to its experimental counterpart. Volume of all the ‘C_lysed’ cell types are subjected to 0.1 unit volume decrease at each MCS to mimic dissipation of degraded matrix materials *in vivo.*

#### Growth and proliferation

Cancer cell*s* are designed to grow in linear combination of two processes.

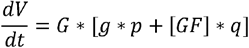

Where V = volume of cancer cell

g = measure of nutrient availability

[GF] = concentration of growth factor (GF) at center of mass of ‘CELL’ p, q = constants

G= Growth rate coefficient

The common surface area of a cancer cell with its neighboring cancer cell (k) and the total cell surface area (s) is accessed to calculate *g* in this equation as *g=(s-k)/40*. The denominator in the calculation of *g* is due to 2D nature of the simulation as a cell can be surrounded by other cell*s* only in xy plane and not in z axis. The scaling of that extra cell surface area without any neighboring cells in z axis is provided by the denominator. Another contributor of the growth function is *[GF]* which mimics the ECM-degradation dependence of growth and proliferation (Olivares et al., 2017). The ‘C_lysed’ cell type is programmed to secrete GF at each of its pixel location where the diffusion constant is kept low (0.02) to localize this growth signal to areas of matrix degradation. p (=1/12) and q (=1/21) constant values are set according to the assumed weightage of the two variables in growth equation. Value of G, then determines how much the resultant growth of the cell should be. The linear combination of the two contributors is multiplied by G, this G is changes during parameter scan. Hence, G or growth rate coefficient is considered as an input variable.

Cell division is incorporated into the cancer cell*s* by a CC3D steppable called ‘MitosisSteppable’ with base function ‘MitosisSteppableBase’. If any cancer cell reaches a threshold volume of 30 units then that cell will be divided in random orientation. The resultant two cells will have volumes half of its predecessor with all other properties kept same as the parent cell. In this model, growth rate is directly correlated to proliferation as it determines the volume of the cell to reach threshold for cell division.

#### Chemotaxis and migration

Until this attribute is added to the cells, the model depended overtly on proliferation for the invasion that was observed and characterized. The absence of a chemoattractant(s) in the framework resulted in this behaviour. We have mitigated this deficiency by incorporating chemoattraction in our revised model. The stromal ECM, collagen has been shown to have chemoattractant properties in undegraded and degraded states (Postlethwaite et al., 1978) (O’Brien et al., 2010). In the github link provided folder with cc3d codes, it can be seen that the .xml file contains “Chemotaxis” plugin for directed chemotaxis induced by Collagen.

The ‘Chemotaxis’ plugin implements a force on the cells through the Hamiltonian towards positive gradient of the chemical field ‘GF’. The ‘GF’ field is secreted by degraded matrix (C_Lysed) and undegraded collagen (C1). In order to ascertain local positive chemical gradient towards undegraded C1 matrix, the C1 releases the chemoattractant in slightly greater amount than its degraded counterpart C_Lysed [secretion rate: 2.5 for C1 compared to 1.0 for C_Lysed]. Only the C1 which are in the vicinity of degradation release this GF as instructed in the ‘SecretionSteppable’. This is coded in the cancol2Steppables.py file in the CC3D folder and ‘Simulation’ subfolder.

#### Parameter scan

CC3D’s parameter scan feature was used to explore parameter space of the input variables. All the combinations of provided values of the parameters are considered for each individual simulation.

Here are the 5 values of the 6 input variables for a parameter scan:

**RD cooperativity**: 2,4,6,8,10

**MMP (A) diffusion constant**: 0.005, 0.01, 0.025, 0.04, 0.055

**Cell-cell contact energy**: 4,18, 32, 46, 60

**Cell-laminin contact energy**: 4,18, 32, 46, 60

**Cell-C1 contact energy**: 4,18, 32, 46, 60

**Cell Growth (G)**: 0, 0.25, 0.5, 0.75, 1

3 replicates of the parameter scan were performed.

The first two values of the input values are considered low values, the 3^rd^ value as intermediate and the last 2 values are considered high values in further analysis.

#### Matlab-based analysis

<u>Quantification of invasion</u>: [Fig1 (B)]

Graphical representation of all the cell*s* in the simulation grid is captured as images at different MCSs for each simulation. A certain ‘timepoint’ or MCS (=1500) is chosen so that MATLAB accesses all the images of all simulations at MCS 1500 and provides quantification. The choice of the 1500^th^ MCS is guided by the fact that this represents the stage where collective invasion can clearly be discerned from the dispersed cells. Beyond this stage, the dispersed cells move out of our framework window and the collectively invasive mass fills the same, disallowing distinctions to be made between invasions of different scales. First, we have binarized the image using the tool ‘Color Thresholder’ (for batch processing, the function from this thresholder tool was passed through ‘Image Batch Processor’). After binarization basic image manipulation functions such as ‘imfill’, ‘imdilate’, ‘bwperim’ were used to optimally isolate the objects from the image. ‘Image Region Analysed was used to get different quantification parameters of those binarized objects. More details can be found from the matlab code (Fig1_simulation_image_analysis.m) provided in github.

##### No. of dispersed objects

The collected simulation images are originally in RGB color scheme to differentiate cell types as displayed in Fig 1(C). Individual images from all simulations are batch-processed to generate binary images with respect to cancer cell type (red color). The first set of binarized images undergo series of rudimentary Matlab image processing operations to generate different set of binary images for different analysis. From them, all the boundaries in each image are then identified to recognize areas which are enclosed by a continuous non-overlapping boundary. These areas without shared boundaries originally represent cancer cell clusters in the respective simulation grid. These clusters can contain single to multiple cancer cell*s*. the number of these clusters denote the dispersiveness of the initial tumor mass in the simulation pertaining to certain combination of input variables, therefore considered as one of the output variables i.e. no. of dispersed objects. The values of this quantity pertaining to each simulation is plotted on the y-axis of the phenospace.

##### Area of largest cell cluster

From the sets of binarized images, areas of all dispersed objects are calculated by counting the pixels preset inside each object’s boundaries. The dispersed object having biggest area is considered the main collective cancer cell cluster of the respective simulation. The value of the area of such as object in the simulation image is quantified as ‘area of largest cell cluster’, the other output variable. The values of this quantity pertaining to each simulation is plotted on the x-axis of the phenospace.

#### K-means clustering

Broad guidelines for performing the Silhouette analysis were adapted from the following link (https://in.mathworks.com/help/stats/kmeans.html#bues3lh). K-means clustering or Lloyd’s algorithm is used for subsequent analysis of the phenospace using MATLAB (Lloyd, 1982). It is an iterative, data-partitioning algorithm that assigns all observations from simulations to exactly one of the K clusters defined by centroids. Being an unsupervised partition-based algorithm, value of K is chosen before the algorithm starts. ‘kmeans’ function in MATLAB uses a two-phase iterative algorithm to minimize the sum of point-to-centroid distances, summed over all k clusters. The first phase uses batch updates, where each iteration consists of reassigning points to their nearest cluster centroid, all at once, followed by recalculation of cluster centroids. In the second phase, points are individually reassigned, if doing so reduces the sum of distances, and cluster centroids are recomputed after each reassignment, by using feedback updates.

The x and y axis values from the phenospace are normalized (z-score) before the use of ‘kmeans’ function to provide same weightage to the axes of during clustering and at the end of the run, all points are assigned a cluster. The points are then colour-coded in the phenospace to differentiate hosting cluster identities.

#### Silhouette analysis

Broad guidelines for performing the Silhouette analysis were adapted from the following link (https://in.mathworks.com/help/stats/silhouette.html#mw9fe10316-522d-4a3a-9cdd-d39baea0c1ce). The silhouette value for each point is a measure of how similar that point is to points within its own cluster, when compared to points in other clusters. The silhouette value ‘S_i_’ for the ith point is defined as

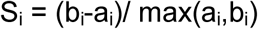

where a_i_ is the average distance from the ith point to the other points in the same cluster as i, and b_i_ is the minimum average distance from the ith point to points in a different cluster, minimized over clusters.

The silhouette value ranges from –1 to 1. A high silhouette value indicates that i is well matched to its own cluster, and poorly matched to other clusters. If most points have a high silhouette value, then the clustering solution is appropriate. If many points have a low or negative silhouette value, then the clustering solution might have too many or too few clusters. So, silhouette values are used as a clustering evaluation criterion for finding a clustering pattern which is invariable across replicates.

After a specific K-means clustering is performed on the points in the phenospace, the normalized matrix of x, y axes along with cluster identity matrix (cluster id of each point) is used to compute silhouette values for each point. The values pertaining to all the points in the simulation is represented in silhouette plot for convenience and the mean of the silhouette values in the plot is denoted with a dashed line. The silhouette method was performed on all the replicates of clustering with a constant k value. The mean of the silhouette values pertaining to each replicate is acquired. The mean and standard deviation of the mean silhouette values associated with replicates of different k values are plotted in a box plot.

#### Principal Component Analysis

Broad guidelines for performing the PCA were adapted from the following link (https://in.mathworks.com/help/stats/pca.html). Briefly, in order to analyse the points in the neighborhoods or patches (denoted by rectangles) in the phenospace, principal component analysis was used in MATLAB. The whole input argument dataset for a cluster’s PCA is a matrix which consists all 6 input variable values for individual simulations relating to the points in that cluster. The matrix is normalized for each variable. Normalization proved to be important as we are interested in relative contribution of the input variable and their changes with respect to other points (simulations), rather than their absolute values. PCA of each rectangle produces different linear combinations of input variables (forms principal component axes) for optimally fitting all points without overlapping in one of those linear relations. PCA quantifies how much of the input data is ‘explained’ by each linear combination (or PC axis) and represented as a scree plot. The linear combination associated with highest y-axis value in the scree plot is identified as first principal component axis. Therefore, the relative contributions of the input variables for constructing PC1 can give insights into the convoluted dynamics of input variables governing the phenotype. The relative contribution of the input variables is plotted in bar graph on the right side of scree plots (Supplementary figure 5).

##### Codes

All the codes relating to the manuscript can be accessed through the following link: https://github.com/drjyprk/cc3d_paper_2_iisc_git

## Acknowledgments

This work was supported by the DBT [BT/PR26526/GET/119/92/2017] SERB[ECR/2015/000280] and the Wellcome Trust/DBT India Alliance Fellowship/Grant [grant number IA/I/17/2/503312] awarded to RB. DP acknowledges KVPY for the scholarship. MKJ was supported by Ramanujan Fellowship awarded by SERB (SB/S2/RJN-049/2018), DST, Government of India.

---

**Supplementary Figure 1:** (A) Initial cell field from the CC3D simulations. (B) The top simulation has Growth rate coefficient set to zero, so, no proliferation property is applied on the red cells. In the bottom image proliferation is active for the cells and they grow to divide as per the growth equation in the materials and methods section. (C) The top simulation has all properties that was used for parameter scan. The bottom simulation did not perform C_Lysed to NC1 conversion, so newly synthesized matrix is not produced after matrix degradation.

**Supplementary Figure 2: (A)** The 3 rows represent similar simulations with 3 varying initial matrix configurations. Left column shows simulation screenshots with only collagen or Collagen matrix (MCS 0, 215, 430). Middle column simulation is with only BM matrix and no Collagen (MCS 0, 2950, 4705). The right column shows the normal setup with BM initially enclosing the cancer cells and that is enclosed by Collagen (MCS 0, 485, 965).**(B)** When the area of minimum enclosing circle is in the range 27000-29200 (bottom image in each column), the simulation screenshots from these 3 simulations were collected and plotted in the phenospace.

**Supplementary Figure 3:** (A) All the simulation outputs in the phenospace (in Figure 1D) are analyzed with K-means clustering using cluster number K=5-8 (leftmost column). Clustering was performed 15 times with random initial cluster centers. The cluster phenotype patterns are shown (middle column) along with their proportional representation within the 15 replicates (right column). (B) The optimal value for K computed using the Silhouette method. Silhouette plots for K=5-8 are constructed with X axis signifying silhouette values and Y axis representing cluster numbers.

**Supplementary Figure 4:** The minimum enclosing circle algorithm used for analysis of invasion in the previous publication was used on the current dataset (Pally et al., 2019). (A) Shows 3 classification of the simulation points in the (K=3)-means cluster-divided phenospace based on the values of the area of the minimum enclosing circle which encloses all the cancer cells in the simulation lattice at 1500MCS. The points having value of this output less than 21852 is colored in green, from 21852 to 43600 is colored blue and more than 43601 is colored red. (B) The statistical plot shows significant difference in the mean of the area of minimum enclosing circle between points of indolent invasion cluster with CMI or DMI clusters. Statistical test was performed using unpaired ANOVA with Tukey’s post hoc multiple comparison (p<0.0001)

**Supplementary Figure 5:** Three principal component analysis data is shown belonging to the individual clusters from k-means clustering with k=3. The top plot in the subfigures show the placement of simulation points of the k(=3)-means cluster in the 3-D space constructed with first 3 principal component axes. In the bottom left, the scree plot shows percentage of explained variability by each principal component axis. On the bottom right, variable coefficient plot shows fraction of each input variable in the first 3 principal component axis linear combinations.

