## Supplementary figures and images for "Interactive dynamics of matrix adhesion and reaction-diffusion predict diverse multiscale strategies of cancer cell invasion"

### Supplementary Figure 1

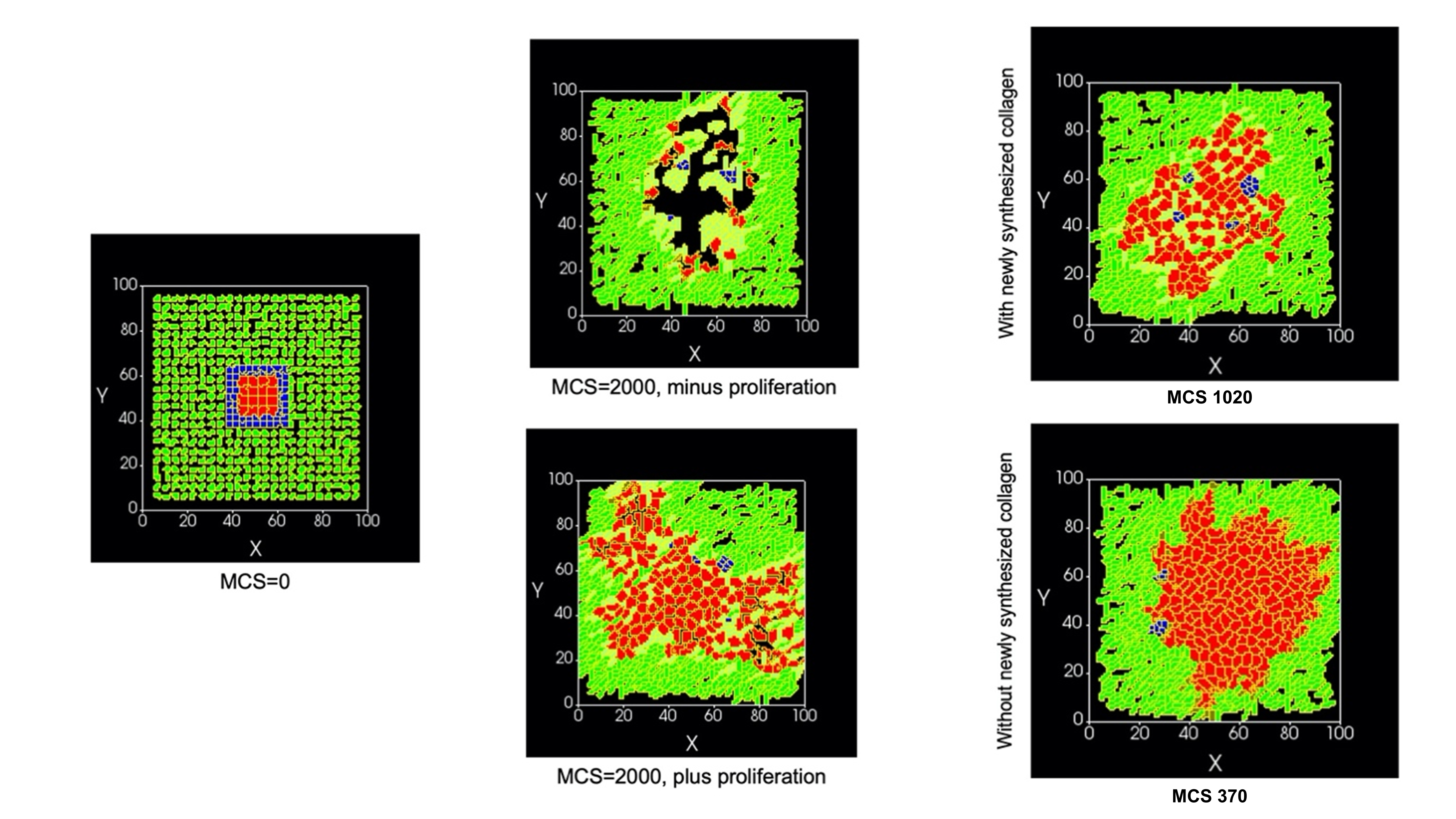

### Supplementary Figure 2

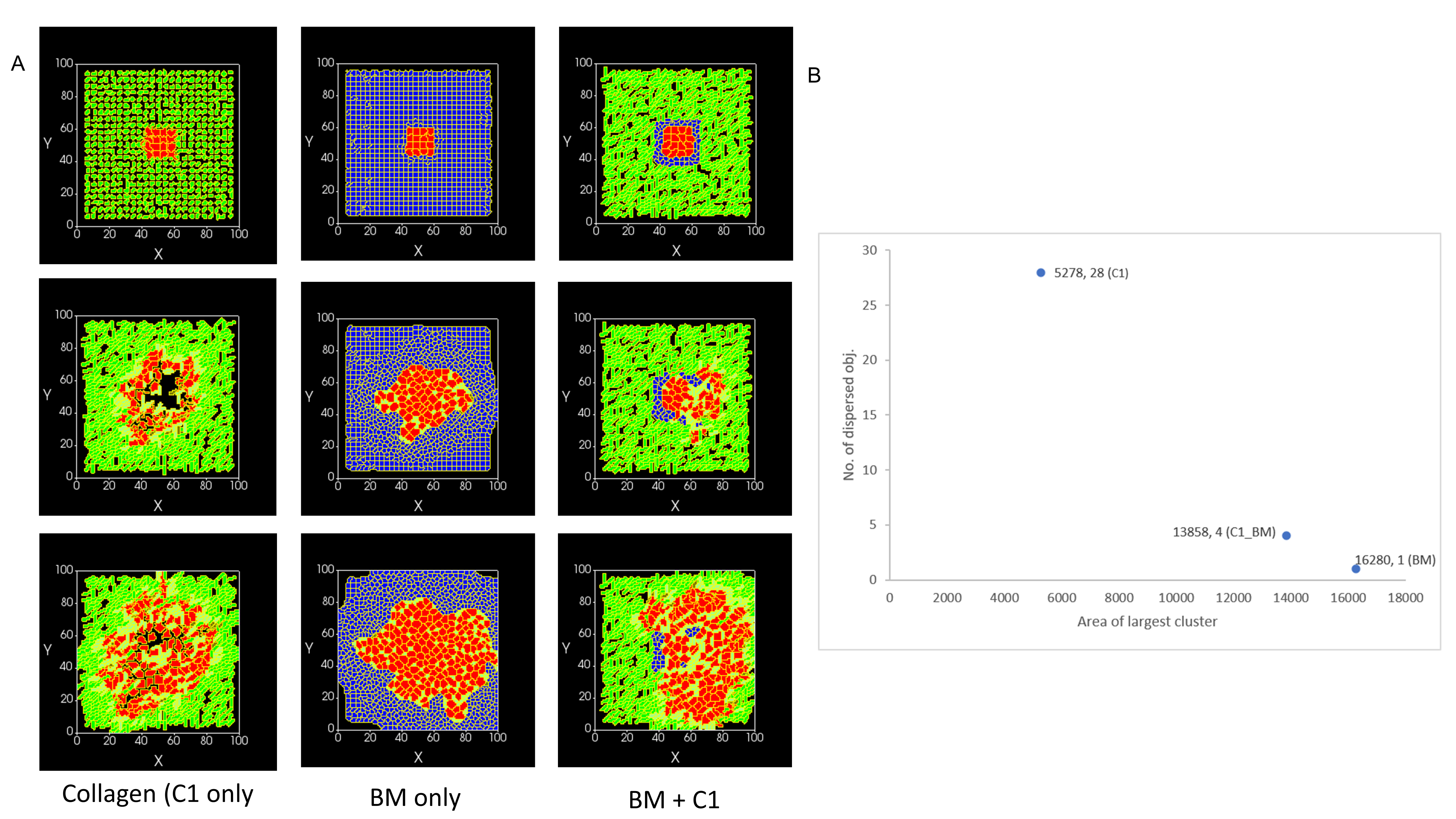

### Supplementary Figure 3

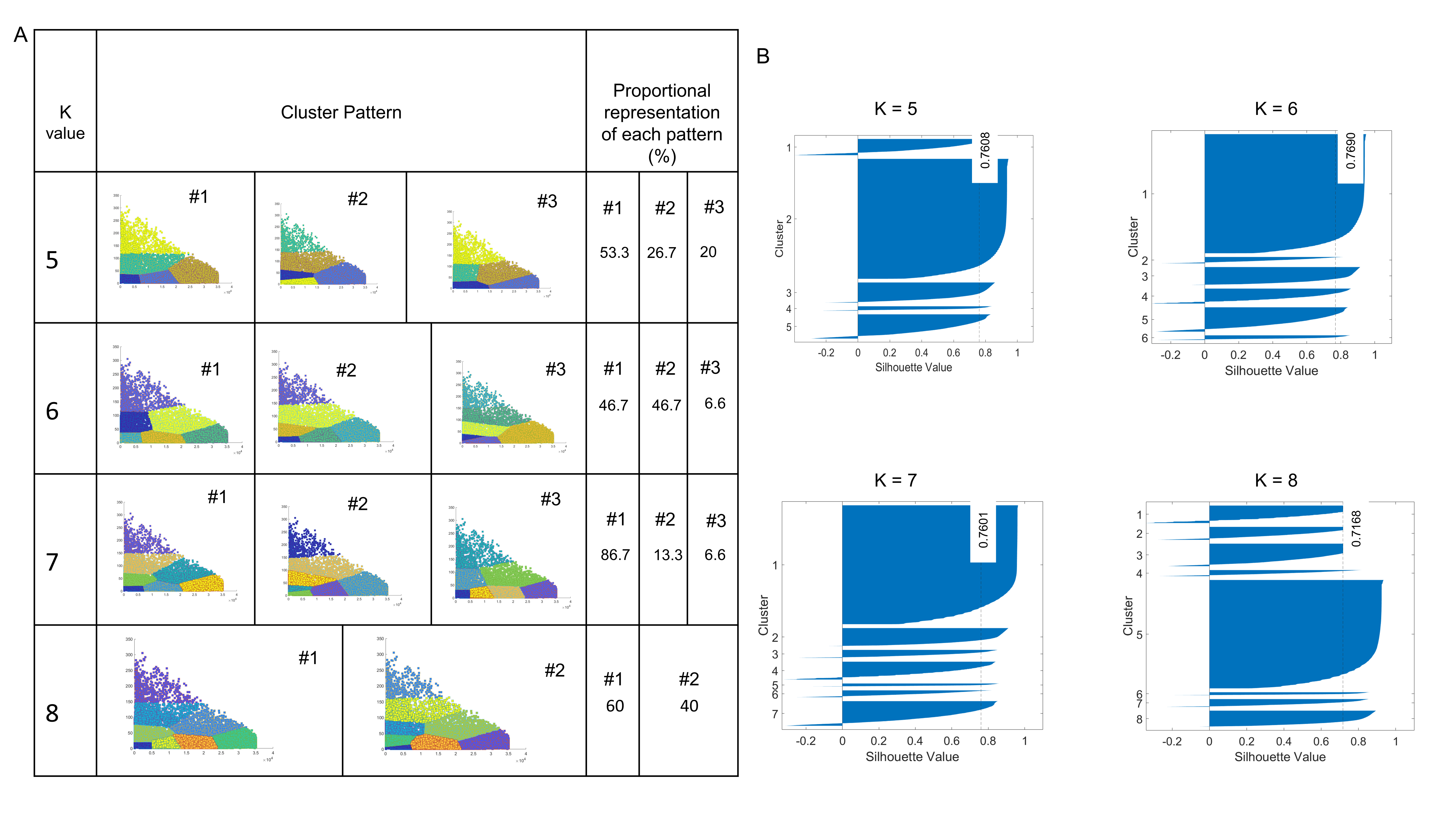

### Supplementary Figure 4

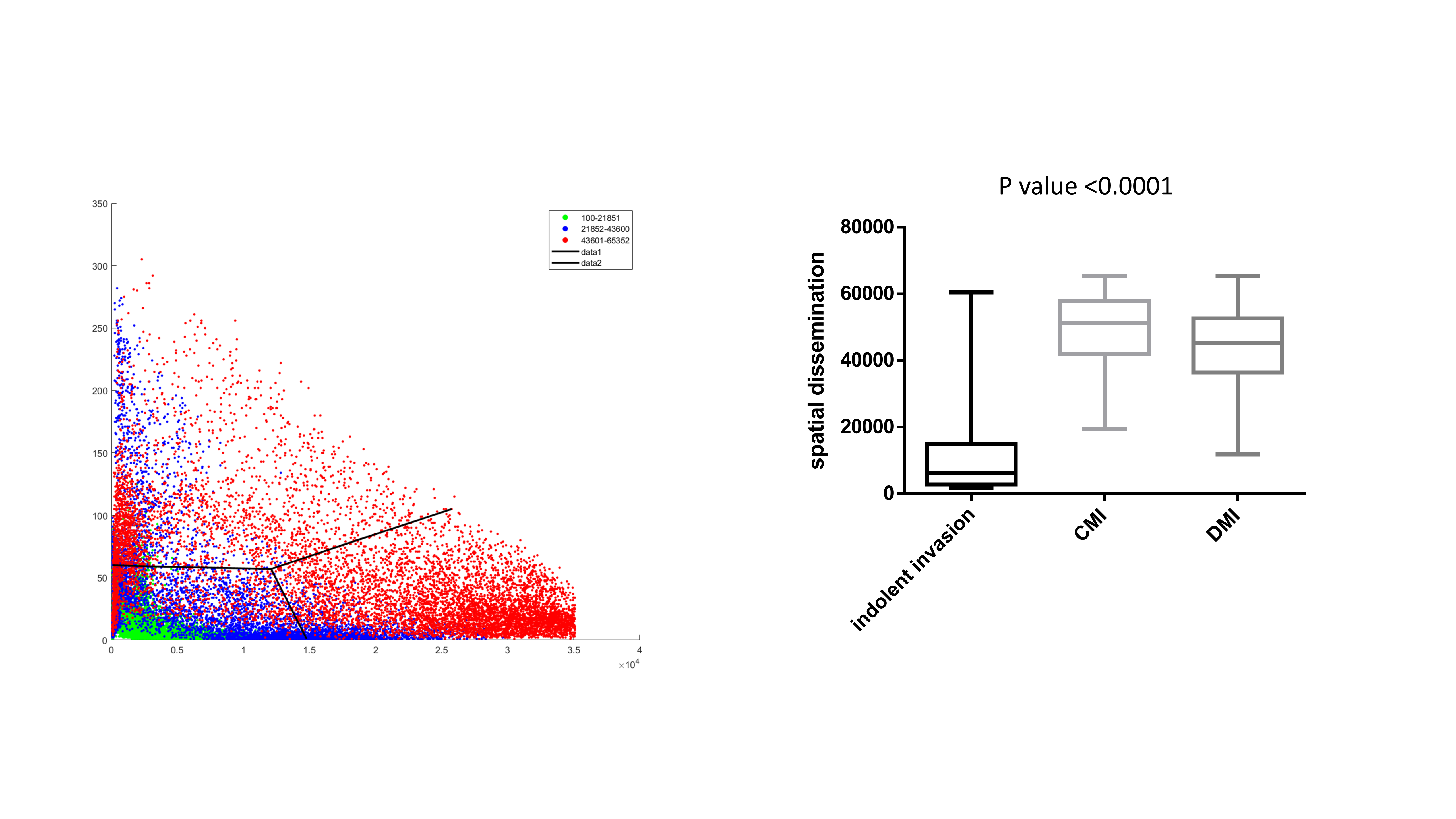

### Supplementary Figure 5

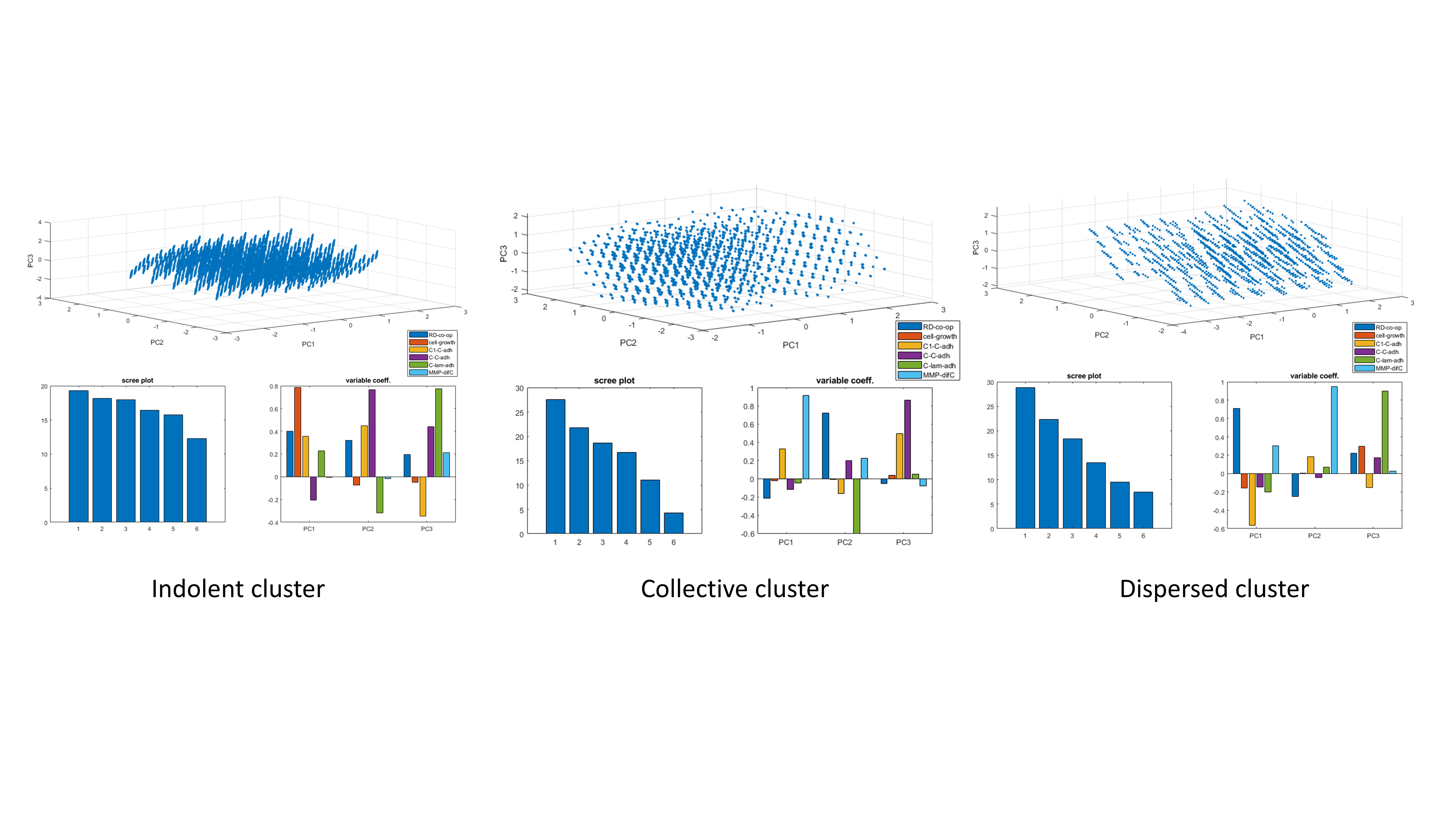
